# CryoLike: A python package for cryo-electron microscopy image-to-structure likelihood calculations

**DOI:** 10.1101/2024.10.18.619077

**Authors:** Wai Shing Tang, Jeff Soules, Aaditya Rangan, Pilar Cossio

## Abstract

Extracting conformational heterogeneity from cryo-electron microscopy (cryo-EM) images is particularly challenging for flexible biomolecules, where traditional 3D classification approaches often fail. Over the past few decades, advancements in experimental and computational techniques have been made to tackle this challenge, especially Bayesian-based approaches that provide physically interpretable insights into cryo-EM heterogeneity. To reduce the computational cost for Bayesian approaches, we introduce CryoLike, a computationally efficient algorithm for evaluating image-to-structure (or image-to-volume) likelihoods across large image datasets, which is built on Fourier-Bessel representations of the images and packaged in a user-friendly Python workflow.

## 1 Introduction

In single-particle cryo-electron microscopy (cryo-EM), an electron microscope captures 2D projection images of individual biomolecules, each in an unknown conformation and pose (orientation and location). Single-particle reconstruction algorithms have been widely used to obtain high-resolution 3D reconstructions [1]. These algorithms use a small subset of images (particles) to generate one or a few 3D maps (volumes) by approximating image-to-volume likelihoods using expectation-maximization [2–4], or high-resolution refinement [5, 6]. The key requirement for high-resolution reconstruction is that the particles within a given subset or class must originate from the same conformation. However, many biologically important molecules exist in multiple conformations rather than a single, static conformation. As a result, isolating particles that represent a single conformation often involves extensive filtering and classification, which can lead to discarding a large portion of particles [7]. These discarded particles might represent rarely occupied conformations or transition states, which can be biologically significant. This limitation has driven development of advanced cryo-EM heterogeneous reconstruction methods [8–16]. However, these approaches still face significant challenges, particularly when no consensus reconstruction can be obtained for highly flexible biomolecules.

In cryo-EM, each particle preserves information about each individual conformation, and in principle, if no particle subsets were discarded, it would be possible to recover the conformational probability distribution of the biomolecules by, *e.g*., using Bayesian approaches [8,17,18], which can be related to the free-energy landscape of the system. However, this requires an accurate evaluation of the image-to-structure likelihoods, which is computationally demanding, especially when the unknown poses are resolved by sampling the likelihood across a large set of possible poses. When performing high-fidelity discrimination of molecular conformations [19, 20] or identifying molecules in crowded environments with cross-correlation-based approaches [21], these computations can be prohibitively expensive for large datasets of structures and images (*e.g*., thousands of structures from molecular dynamics and millions of particles from a typical cryo-EM experiment). Therefore, efficient algorithms for calculating the image likelihood for cryo-EM will significantly reduce the computational overhead of these methods.

Previously, Barnett *et al*. [22], Rangan *et al*. [23], and Rangan *et al*. [24] have presented a rapid and mathematically efficient algorithm for *ab initio* reconstruction using Fourier-Bessel representations to compute the cross-correlation between cryo-EM images and 2D templates (*i.e*., projection images of the 3D structure). Based on the mathematical foundation built by these works, we develop “CryoLike” (https://github.com/flatironinstitute/CryoLike), a GPU-accelerated cryo-EM software designed to compute structure-to-image likelihoods. CryoLike implements a computationally efficient algorithm for evaluating the image-to-structure likelihood in cryo-EM, provided within a user-friendly Python workflow. In this paper, we first detail the architecture of the CryoLike software and explain the mathematical foundations underlying template generation, cross-correlation, and likelihood computations (described in detail in the Appendix). Then, we present computational benchmarks to demonstrate the performance of CryoLike and showcase its applicability in model discrimination using experimental cryo-EM data for three different proteins: apoferritin, hemagglutinin, and the SARS-CoV-2 spike protein. We finalize with the conclusions of the work.

## 2 Methods

### 2.1 Software architecture

Fig. 1(a) provides an overview of CryoLike’s software infrastructure. CryoLike takes the following inputs: (*i*) particle stacks in MRC format, (*ii*) CTF parameters in STAR or CryoSparc format, and (*iii*) either atomic coordinates in PDB format or a 3D volume in MRC format. The following parameters are optional inputs (default values, shown in Appendix Table 1, are used if not provided): (1) a resolution factor, which defines the maximum resolution for the images and modified projections of the volume (*i.e*., templates, see section 2.2); (2) the maximum 2D displacement in pixels; (3) the total number of displacements; and (4) the atomic models, with the option to use the Van der Waals model on the *C*_*α*_ [20] or to specify atomic radii for each atom, and (5) the angular distance between viewing angles of the templates, *i.e*., the sampling density of templates on a unit sphere of the viewing angles, and (6) the number of in-plane angles sampling the Fourier densities for the images and templates.

**Table 1:** Summary of mathematical symbols.

| Symbol | Description |
| --- | --- |
| $\mathbf{x}$ | 3D coordinates in physical space |
| $\mathbf{k}$ | 3D coordinates in Fourier space |
| $F(\mathbf{x})$ | 3D cryo-EM volume in physical space |
| $\hat{F}(\mathbf{k})$ | 3D volume in Fourier space |
| $\beta, \alpha$ | Viewing direction defined by a polar tilt angle $\beta$, and azimuthal angle $\alpha$ |
| $\hat{S}l \circ \mathcal{R}(\beta, \alpha)$ | Projection operator defined by $\beta$ and $\alpha$ |
| $\delta, \gamma$ | 2D displacement $\delta$ and inplane rotation angle $\gamma$ |
| $\mathcal{R}(\gamma), \mathcal{R}(\beta, \alpha)$ | 2D rotation operator defined by $\gamma$ and 3D rotation defined by $\beta$ and $\alpha$ |
| $\mathcal{T}(\delta), \hat{\mathcal{T}}(\delta, \mathbf{k})$ | Translation operator in physical and Fourier space defined by displacement $\delta$ |
| $\tau = \{\gamma, \beta, \alpha, \delta\}$ | Image parameters including viewing angles, 2D displacement, 2D rotation |
| $\mathbf{K}(\beta, \alpha)$ | Fourier slice in 3D Fourier space defined by $\{\beta, \alpha\}$ |
| $\mathbf{k}_{\text{polar}} := (k, \psi)$ | 2D polar coordinates in Fourier space describing the projected 2D space $\mathbf{K}$ |
| $S(\mathbf{x}; \tau, \text{CTF}, F)$ or $\hat{S}(\mathbf{k}; \tau, \text{CTF}, \hat{F})$ | 2D template: projection of the model $F$ at a particular $\tau$ with CTF applied |
| $A, \hat{A}$ | 2D image in physical space and Fourier space |
| $n_{\text{p}}, N_{\text{pix}}$ | number of pixels along one axis and the total number of pixels in the image |
| † | Complex conjugate |
| $R_i, \mathbf{r}_i$ | Atomic radius $R_i$ and atomic positions $\mathbf{r}_i$ of atom $i$ |
| $\nu, \mu$ | Image-specific intensity factor $\nu$ and offset $\mu$ |
| $\varepsilon(\mathbf{x}), \lambda$ | Pixel Gaussian white-noise model with standard deviation $\lambda$ |

**Figure 1:**
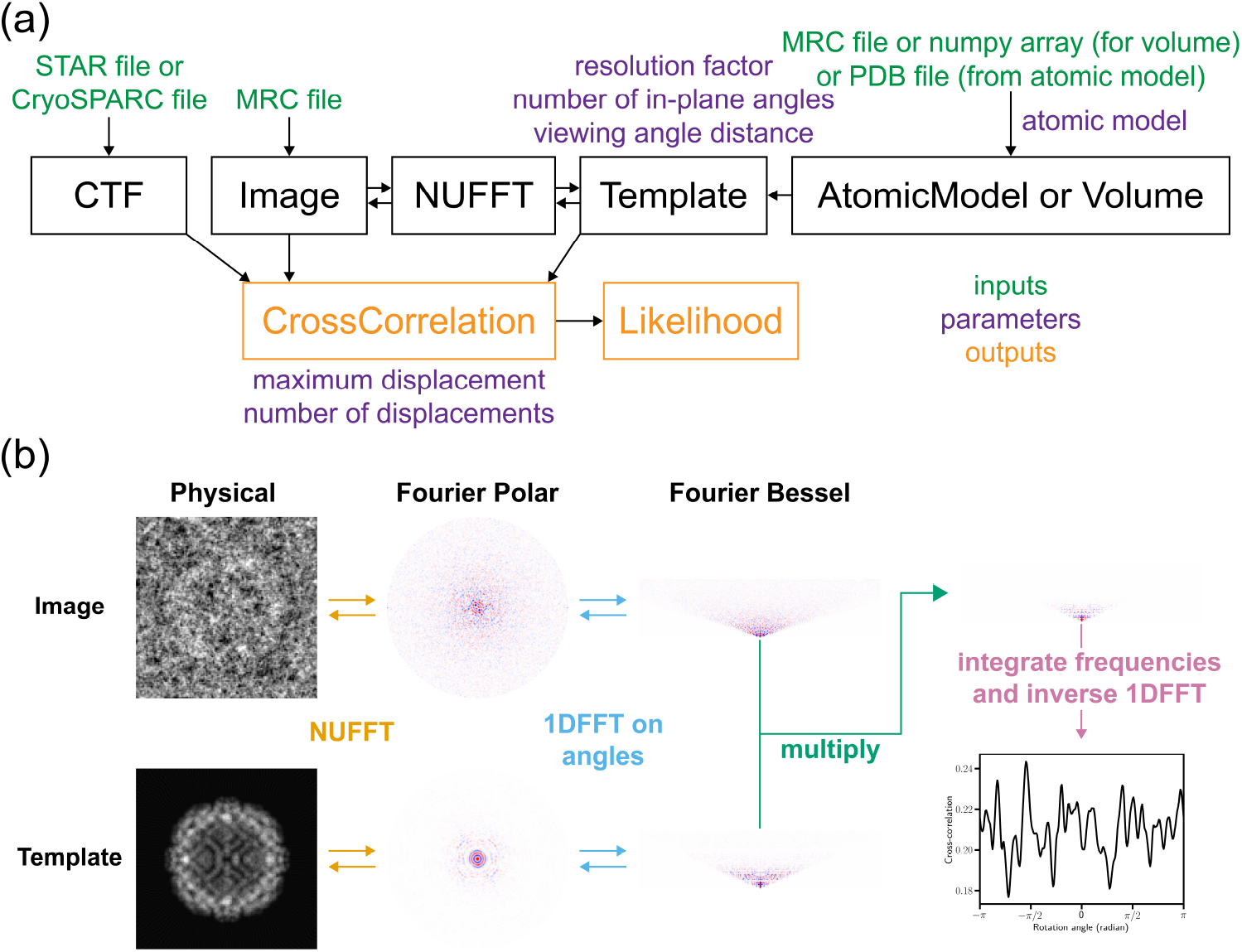
Overview of the Cryolike software pipeline. **(a)** Workflow diagram: The input images are read from a STAR or CryoSPARC file (containing the metadata) and an MRC file (containing the image data). A non-uniform fast Fourier transform (NUFFT) converts the images from physical space to Fourier space. The sampling density of the images and templates is determined by the resolution factor and viewing angle distance. A set of templates is derived from an atomic model (from a PDB file) or volume data (from an MRC file). The contrast transfer function (CTF) (corresponding to each image) is applied to the templates during the cross-correlation calculation. The cross-correlation values between the images and the templates at all possible image parameters *τ* are calculated. The outputs of CryoLike are the optimal cross-correlation, optimal parameters *τ* that maximize the cross-correlation, and the likelihood values for each image. **(b)** Image transformation: The physical image and template are transformed via NUFFT between physical space represented by cartesian coordinates and Fourier space represented in polar coordinates. The Fourier polar images undergo a 1D Fast Fourier Transform (1DFFT) across the polar angular axis, giving the Fourier-Bessel coefficients. The Fourier-Bessel coefficients of an image and a template are multiplied, then integrated across the frequencies, and then inversed 1DFFT back to the Fourier domain, resulting in the cross-correlation with respect to the rotation angle (bottom right).

CryoLike generates templates from a structure or a volume, converts the particle stacks into Fourier images, calculates the cross-correlation between the images and templates (Fig. 1(b)), and computes the image-to-structure likelihood. CryoLike generates output files containing the cross-correlation at the optimal pose between each image and structure (or

volume), the optimal pose for each image (including polar and azimuthal viewing angles {*β, α*}, 2D image displacement ***δ***, and in-plane rotation *γ*), and the log-likelihood of each image given the 3D structure. Table 1 summarizes the relevant mathematical notation.

### 2.2 Template generation

According to the Fourier-slice theorem, the Fourier Transform of a 2D projection of the physical density *F* (**x**) is equivalent to the Fourier density 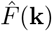 evaluated on the set of frequencies **k** ∈ **K**(*β, α*) ⊂ ℝ^3^, where **K**(*β, α*) is a ‘2D slice’ that passes through the origin of ℝ^3^ with its orientation defined by the viewing angles {*β, α*}. Here, *β* is the polar tilt angle, and *α* is the azimuthal rotation angle of the projection. CryoLike samples the viewing angles 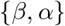 by a discrete set of *n*_template_ roughly equispaced points on a unit sphere. We define a projection operator 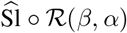 that extracts this Fourier slice from 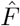, *i.e*., 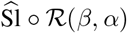 evaluates the Fourier density 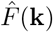 for all points **k** on **K**(*β, α*). In practice, we sample 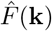 on a discrete polar grid {***k***_polar_} := {(*k, ψ*)} ∈ ℝ^2^ parametrizing **K**(*β, α*), where the radii {*k*} are defined by a Gauss-Jacobi quadrature with *n*_*k*_ nodes on the interval [0, *k*_max_], with *k*_max_ the maximum frequency (*i.e*., the Nyquist sampling-rate when we sample *F* to full resolution), and the angles {*ψ*} are defined by a discrete set of *n*_*ψ*_ equispaced points on [0, 2*π*), where *n*_*ψ*_ is the number of in-plane angles.

CryoLike takes an atomic model or a 3D volume as input and generates templates in Fourier space. When given the PDB model, the cryo-EM density of each atom is represented as either a 3D Gaussian distribution (g) or a hard sphere (h). We evaluate the template intensity at each **k** ∈ ℝ^3^ by

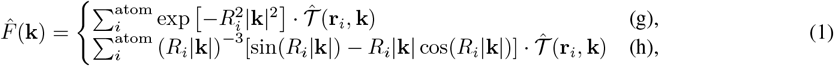

where 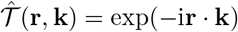 is the 3D translation kernel in Fourier-space, and *R*_*i*_ and **r**_*i*_ are the atomic radius and position of the *i*^th^ atom, respectively. We use the hard-sphere model with Van der Waals radii on the *C*_*α*_ atoms for the results shown in this paper. When given a real space 3D density *F*(x) sampled at a collection of spatial points **x** ∈ ℝ^3^, we evaluate 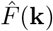 at each required **k** ∈ ℝ^3^ by using a 3D non-uniform fast-Fourier-transform (NUFFT). In both cases, each Fourier slice 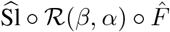 then corresponds to the Fourier transform of a projection of the original volume *F* along the viewing angles {*β, α*}. These slices can then be (i) further rotated by an in-plane rotation *γ*, (ii) displaced by a 2D translation ***δ*** (via multiplication by the 2d translation kernel 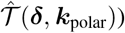 and (ii) transformed via an image-specific contrast transfer function, or ‘CTF’ to produce the associated templates 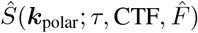,where *τ* := {*γ, β, α*, ***δ***} is the collection of pose-parameters which can determine a template, and we will refer to 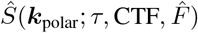 as 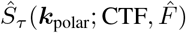 or 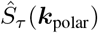 when the context is clear. The collection of all such templates 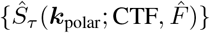 will be used below to assess the likelihood of generating any particular image given the volume or structure 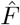.

### 2.3 Image preprocessing

The intensities of the input physical images are centered at 0. The physical images are then transformed via a 2D NUFFT into Fourier images *Â*(***k***_polar_) using the finufft package [25–27]. The total power in Fourier space is normalized to 1 for the templates and images. For example, 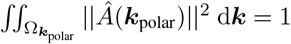, where 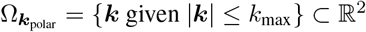 is the frequency-domain on which the images are resolved. An example of this transformation is shown in Fig. 1(b).

### 2.4 Cross-correlation computation

In its most basic form, the cross-correlation *χ* (*γ, f, g*) between two functions *f* (*ψ*) and *g*(*ψ*) is defined as

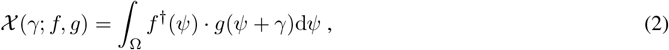

with ^*†*^ denoting complex-conjugation and Ω denoting the domain of integration. The cross-correlation can be evaluated using the convolution theorem by taking the product of the Fourier transforms

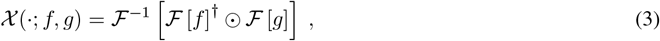

where ℱ denotes the forward 1D Fourier transform, and element-wise multiplication is denoted by ‘⊙‘. CryoLike applies a convolution theorem of this form to the angular coordinate in 2D Fourier space. More specifically, given any image *Â*(*k, ψ*) and template 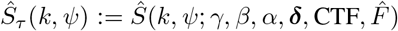, CryoLike calculates the frequency-space cross-correlation *χ* _freq_(*γ*; Ŝ, *Â*) by combining a radial integration with an angular convolution. In practice, as shown in Fig. 1(b), CryoLike first performs a 1D Fourier transform along the angular coordinate *ψ* to convert the Fourier images and templates into a Fourier-Bessel basis, then multiplies the Fourier-Bessel coefficients of the image and the template, and finally transforms the multiplied coefficients back to the original Fourier basis to obtain the cross-correlation in Fourier space with respect to each 2D rotation *γ*. Formally, this cross-correlation is given by

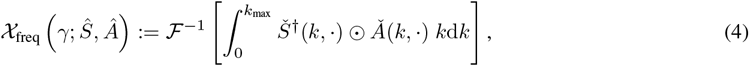

where 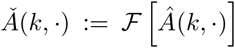 and *Š*(*k*, ·) := ℱ [*Ŝ*(*k*, ·)] are the Fourier-Bessel coordinates associated with the template and image, respectively. The cross-correlation in Eq. 4 is a function of the image *Â*, its CTF, and the 3D volume 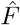,as well as the various pose parameters *τ* := {*γ, β, α*, ***δ***}. For brevity we will refer to this cross-correlation as 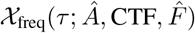.CryoLike calculates the cross-correlation for a discrete collection of *τ*. We can then use this to select the optimal image pose *τ* ^opt^ for each image *Â* by

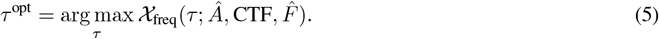

This optimal pose is then used to calculate the cross-correlation between the corresponding template and the image in physical space, producing 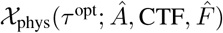 as described in the Appendix. Note that, while often very similar, *χ* _freq_ and *χ*_phys_ will not be exactly the same because our domain of integration 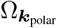 is circular in shape, excluding the ‘corners’ of the cartesian frequency grid typically associated with the array of pixels in real space.

### 2.5 Image-to-2D-template likelihood

In this section, we present a simplified formulation of the image-to-template likelihood in physical space, which is easily transferable to Fourier space. The full derivation of both the physical and Fourier image likelihood is presented in the Appendix. In physical image space, we assume a Gaussian white-noise model *ε*(***x***). We express an image *A*(***x***) in terms of a particular projection *S*_*τ*_ (***x***) at pose *τ*, multiplied by an image-specific intensity ν and added to an offset *µ*,

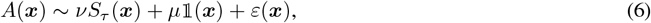

where 𝟙(***x***) denotes the support of the real-space pixel-grid. Here, we assume *ε*(***x***) is a Gaussian white noise model, *i.e*., each pixel-value of *ε*(***x***) is drawn independently from the distribution 𝒩 (0, *λ*^2^), where the pixel variance *λ*^2^ is constant across all frequencies.

The basic likelihood of observing the image *A* given a particular template *S*_*τ*_ is then

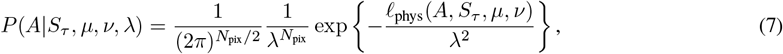

with *ℓ*_phys_(*A, S*_*τ*_, *µ, ν*) denoting the l2-norm of the difference between the image *A* and template *S*, given the intensity ν and offset *µ, i.e*.,

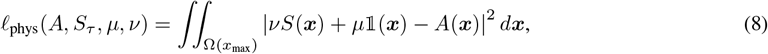

with Ω(*x*_max_) corresponding to the physical domain of integration. We detail the expression of *ℓ*_phys_ and its Fourier counterpart *ℓ*_freq_ in terms of the cross-correlations *χ*_phys_ and *χ*_freq_, respectively, in the Appendix.

### 2.6 Image-to-structure likelihood: marginalization over parameters

We assume a flat prior over all parameters. We marginalize the image-to-2D-template likelihood (Eq. 7) over image-specific intensity factor ν, offset *µ*, and the pixel variance *λ*^2^ analytically, similarly as in ref. [19]. We detail this derivation of the marginalized likelihood in the Appendix. After the analytical integration, CryoLike provides three different ways to approximate the likelihood *P* (*A*| *S*_*τ*_) of an image, given a model: (i) evaluating *P*(*A* | *S*_*τ*_) in physical space for only *τ* ^opt^ (“optimal physical”); (ii) evaluating *P*(*Â*| *Ŝ*_*τ*_) in Fourier space for only *τ* ^opt^ (“optimal Fourier”); and (iii) evaluating *P*(*Â* | *Ŝ*_*τ*_) in Fourier space for all possible templates *Ŝ*_*τ*_ and then integrating *P*(*Â* | *Ŝ*_*τ*_) over *τ* (henceforth “integrated Fourier”), *i.e*.,

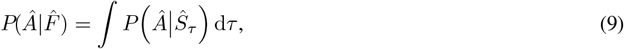

where we have assumed a uniform prior over *τ*. We do not incorporate the physical-space equivalent of (iii) (“Integrated physical”), as this calculation requires evaluating *P* (*A* | *S*_*τ*_) for templates *S*_*τ*_ in physical space for all possible *τ*, which requires extra steps to compute.

## 3 Results

### 2.1 Computational performance

The cross-correlation is the most computationally costly part of the algorithm. In theory, it has a computational complexity of 𝒪 (*n*_images_*n*_{*β,α*}_*n*_*δ*_*n*_*k*_*n*_*ψ*_), where *n*_{*β,α*}_ is the number of viewing angles for generating templates *S*_{*β,α*}_, *n*_*δ*_ is the number of possible translations, *n*_*k*_ and *n*_*ψ*_ are the number of frequencies and in-plane angles, respectively, in the Fourier polar space. However, for GPU computations, the computational performance heavily depends on the memory management, *i.e*., the batch size of the templates and images being fed into the GPU memory. Therefore, we show the benchmarking results with respect to the batching parameters (*i.e*., the number of templates per batch and the number of images per batch) in Fig. 2(a). The wall clock time decreases as the batch size fed into the GPU increases, indicating that larger batch sizes improve processing efficiency, reducing total computation time. However, when the GPU memory utilization nears its limit at larger batch sizes (*e.g*., with 1024 templates and 1024 images per batch), this performance gain is reduced, likely due to limited available memory.

**Figure 2:**
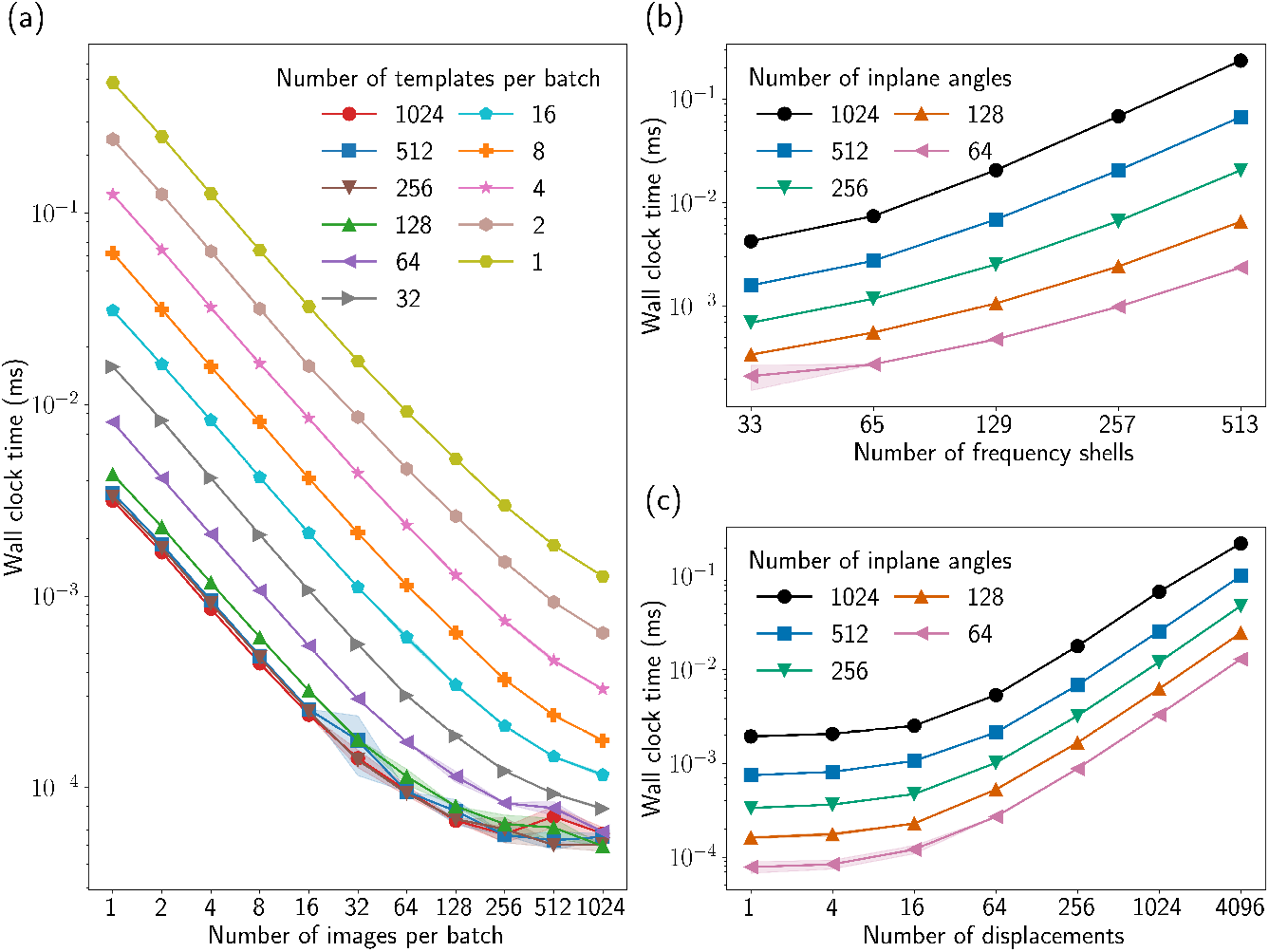
CryoLike wall clock time per image-template pair in milliseconds (ms). **(a)** We show the benchmark wall clock time when using different number of images and templates per batch, *i.e*., the GPU memory batching parameters. **(b)** We show the benchmark wall clock time when using different number of frequency shells and in-plane angles, *i.e*., the dimension of the Fourier polar grid ***k*. (c)** We show the benchmark wall clock time when using differentnumber of displacements and in-plane angles, *i.e*., 2D displacements and rotations. The CryoLike parameters used for the benchmarks are listed in Appendix Table 2. The wall clock time is calculated with one GPU Nvidia H100.

In Fig. 2(b) and (c), we present benchmarking results with respect to the number of in-plane angles, the number of frequency shells, and the number of displacements, respectively. The wall clock time increases as the number of frequency shells grows, showing that the computational cost scales as the resolution increases. Similarly, the wall clock time increases with the number of displacements, showing that the computational cost scales with the size of the search space for optimal 2D alignment.

While the computational performance can vary drastically for hardware setups, the processing time can be estimated by referencing Fig. 2, multiplying the benchmark value by the approximate number of viewing angles (typically around 2000 for standard resolution) and the number of particles to be analyzed. For reference, with an H100 GPU, the comparison of one volume to a particle stack with 1024 images of box size 256 × 256 pixels, with 32× 32=1024 displacement samples, finishes in under 45 seconds, and the same set with box size 128 × 128 pixels, with 16× 16=256 displacement samples, finishes in under 3 seconds.

### 3.2 Model Discrimination and Beyond

To evaluate the efficacy of the likelihood calculation and demonstrate a few real-world use cases, we devised some tests to evaluate the ability of the likelihood to differentiate between a true model and a false model. We computed the log-likelihood ratio (LLR) between a “true model” and a “false model” for each image given by

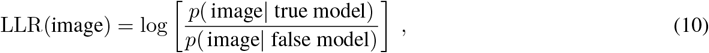

where the true model represents the correct model and the false model is a deformed version of the true model. We use the false negative rate (FNR), *i.e*., the proportion of images with LLR *<* 0, to identify how often the method incorrectly identifies the false model as true. Note that because we are working with real cryo-EM images convoluted with experimental noise, we expect some images to have an LLR *<* 0, *i.e*., false negatives exist.

In Fig. 3, we show the LLR calculated using the “integrated Fourier” likelihood as a function of the images sorted by LLR for three experimental datasets: Apoferritin [28], hemagglutinin [29], and the SARS-CoV-2 Spike protein. For apoferritin and hemagglutinin (Fig. 3(a) and (b), respectively), we use PDB structures 4v1w and6wxb, respectively, as the true model and introduce a deformed version of the structure as the false model (see Appendix Fig. 1 as an example template), which was generated using normal mode analysis. The root-mean-squared deviation of the C_*α*_ atoms between the deformed model and the true model was 7.6 Å and 14.4 Å for apoferritin and hemagglutinin, respectively. We used the 483 particles of apoferritin available from the EMPIAR-10026 dataset and 10000 randomly picked particles of hemagglutinin from EMPIAR-10532. The hemagglutinin dataset contains particles that are not used to generate the high-resolution reconstruction of the true model, which means some particles can deviate from the true model. The CryoLike parameters used are listed in Appendix Table 2. We present similar results calculated using “optimal physical” and “optimal Fourier” in the Appendix Fig. 2. For these two datasets, we compare the CryoLike results with BioEM [19, 20], which uses a different image formation model and likelihood framework, effectively an “integrated physical” that integrates the likelihood over image parameters *τ* in physical space. CryoLike shows an overall improvement in accuracy over BioEM, indicated by the higher LLR for most images and the lower FNR for both apoferritin and hemagglutinin. Due to the older GPU implementation in BioEM, a fair comparison of CryoLike’s computational performance with BioEM is not feasible, as BioEM must operate at a much lower resolution to reach comparable computational speeds.

**Figure 3:**
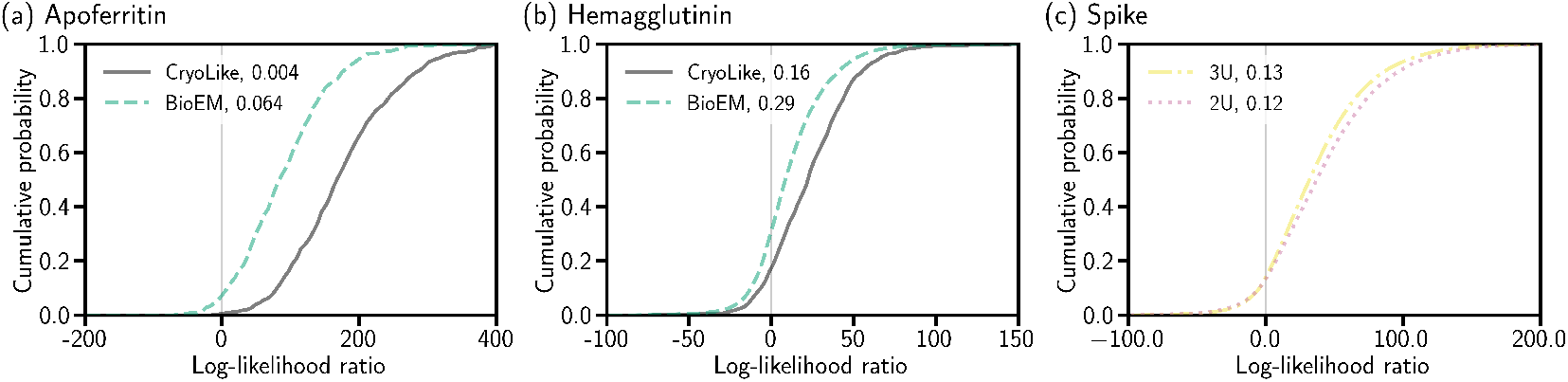
The cumulative probability of the log-likelihood ratio (LLR) between a ‘true model’ and a ‘false model’ (Eq. 10) as a function of the LLR for each image, for (a) apoferritin, (b) hemagglutinin, and (c) two spike protein datasets (3U and 2U), respectively. The CryoLike results are compared to BioEM for (a) and (b). The False negative rate (FNR) is presented after the comma of each legened. See the Main Text for details about the models and image sets.

For the SARS-CoV-2 spike protein, we used cryo-EM images from EMPIAR-12098 [30] for two distinct particle sets: those used to reconstruct EMD-50421 (three RBD up, or “3U”) and those used to reconstruct EMD-50422 (two RBD up and one down, or “2U”). Instead of using atomic structures, we used cryo-EM density maps for two different conformations of the spike protein. Note that the ability to use density maps is a feature that BioEM does not have. Therefore, BioEM is not included in this test. When analyzing the particles for “3U”, we treat “3U” as the true model and “2U” as the false model, and vice versa. For these two cases, the LLR (Fig. 3(c)) is positive for most particles and the FNR is low. To demonstrate the applicability of CryoLike beyond model comparison, we perform ensemble reweighting [17] on the Spike models using the likelihood calculated by CryoLike. We present this result in the Appendix Fig. 3, demonstrating the potential of using the CryoLike likelihood in downstream tasks, such as for estimating conformational probability distribution.

## 4 Conclusions

In this work, we introduced CryoLike, a computationally efficient algorithm designed to evaluate image-to-structure (or volume) likelihoods across large-scale image and structure datasets wrapped in a user-friendly Python workflow.

CryoLike shows exceptional computational performance in evaluating the likelihood and its applicability in model comparison, building on Fourier-Bessel representations. We highlight CryoLike’s potential to be combined with ensemble reweighting techniques [17] to study flexible biomolecules, where conventional cryo-EM heterogeneity methods may fall short due to their reliance on consensus reconstructions.

Future developments include integrating advanced techniques, *e.g*., using the factorization approach [23] to enhance the efficiency of 2D displacement searches, combing it with amortized template matching approaches [31] or using gradient-based optimization for some parameter searches. We also plan to optimize the algorithm for various GPU architectures, particularly for lower-end GPUs with limited internal memory. As we continue to refine and extend the capabilities of CryoLike, we anticipate that it will become an indispensable tool for analyzing dynamic and heterogeneous biological systems with cryo-EM.

## Code Availability

The code is available at GitHub https://github.com/flatironinstitute/CryoLike.

## Acknowledgements

The Flatiron Insititute is a division of the Simons Foundation. The conceptualization of this work was led by A.R., W.T., and P.C. Data curation was carried out by W.T. and P.C. Formal analysis was performed by W.T. Investigation was conducted by W.T. Methodology was developed by A.R., W.T., and P.C. Project administration was provided by P.C. Software was developed by W.T. and J.S. Supervision was provided by P.C. Validation was performed by W.T. Visualization was done by W.T. The original draft was written by W.T. and A.R. Review and editing were provided by W.T., A.R, P.C., and J.S. Computational resources for this work are provided by the Scientific Computing Core and Rusty, a supercomputing cluster at the Flatiron Institute. We acknowledge Erik Thiede for insightful discussions, valuable comments, and thorough reviews of the manuscript and code. We acknowledge Miro Astore for beta-testing the software and providing feedback.

## Appendix Figures

**Appendix Figure 1:**
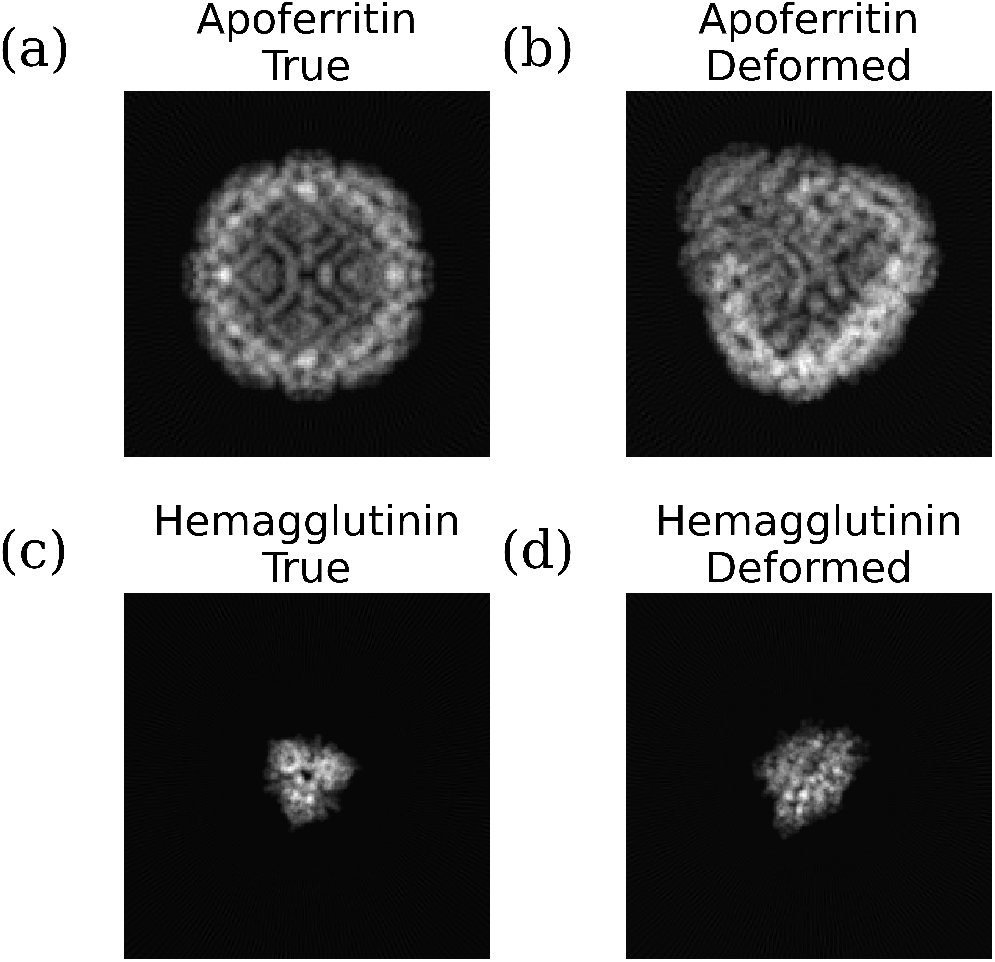
Example templates generated from (a) the true model and (b) the deformed model of apoferritin; and (c) the true model and (d) the deformed model of hemagglutinin.

**Appendix Figure 2:**
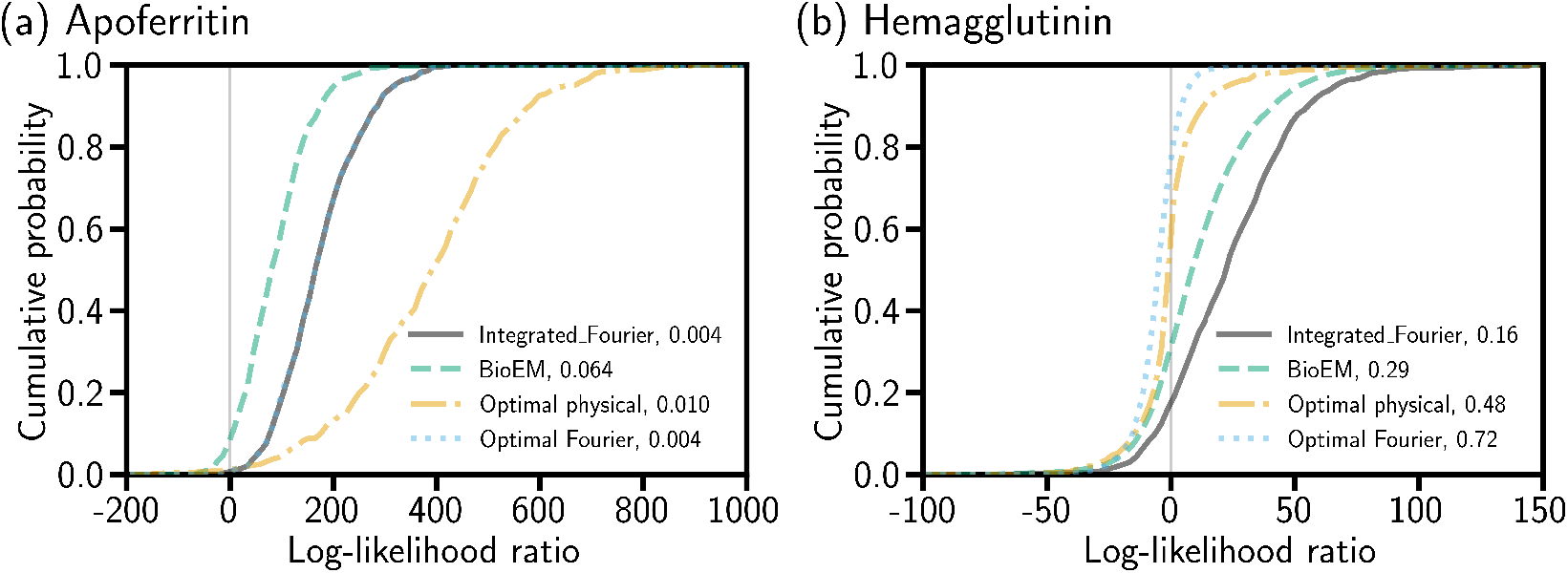
The cumulative probability of the log-likelihood ratio (LLR) between a ‘true model’ and a ‘false model’ (Eq. 10) as a function of the LLR for each image, for (a) apoferritin, and (b) hemagglutinin, respectively. The legend shows the different likelihood methods used to calculate the likelihood with CryoLike and the FNR. “Integrated Fourier” and “BioEM” are as in Fig. 3.

**Appendix Figure 3:**
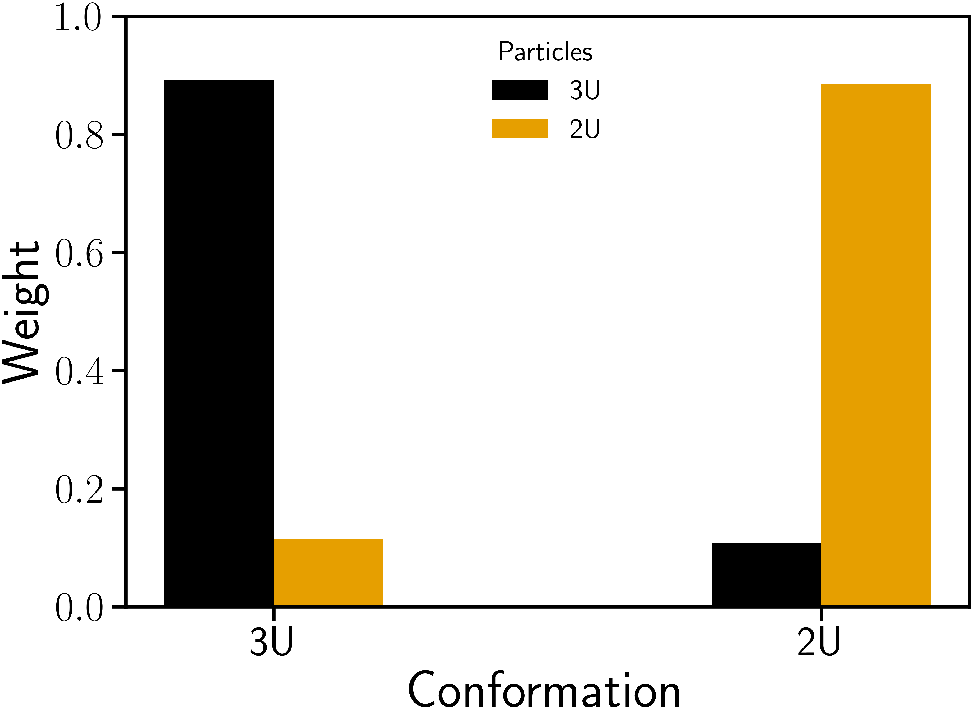
Probability weights computed using the Ensemble reweighting method [17] with the CryoLike likelihood (shown in Fig. 3(c)).

**Appendix Figure 4:**
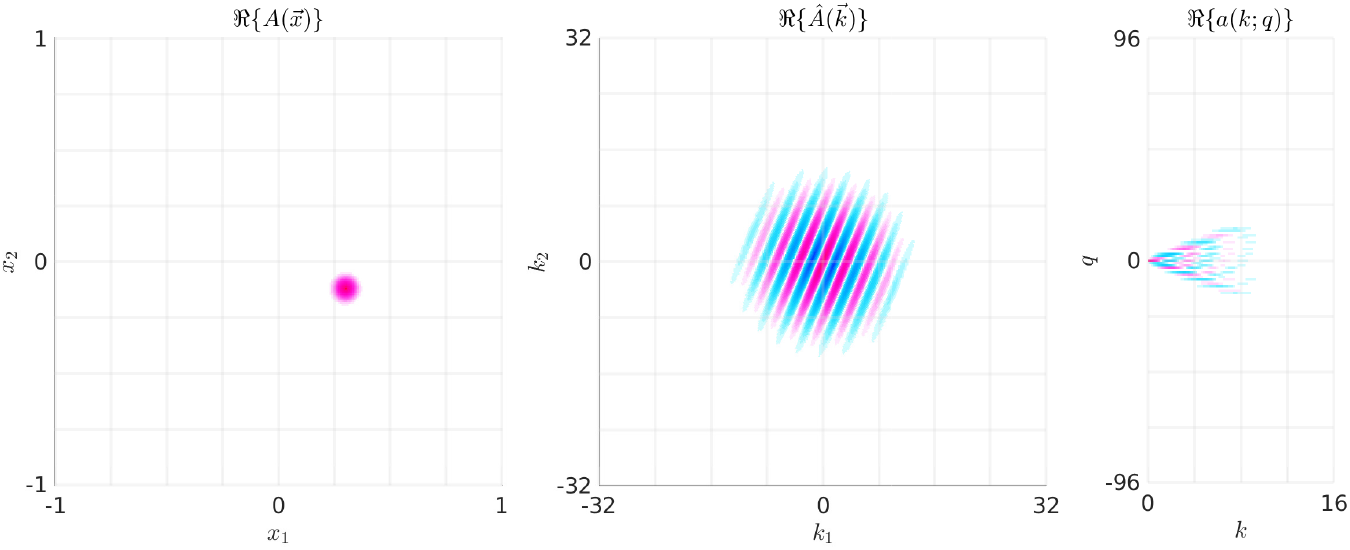
On the left, we show an image *A*(***x***), its Fourier-transform *Â*(*k, ψ*) in the middle, and its Fourier–Bessel representation 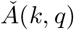 on the right. Each picture depicts a heat map of the real part of the corresponding function, with the colors pink and blue corresponding to positive and negative values, respectively. This image has been chosen so that *A*(***x***) is an isotropic Gaussian in real space, with a mean given by ***x***_0_ = [+0.3, −0.12]. Thus, *A*(***x***) is a mollified version of *δ*(***x*** − ***x***_0_). Consequently, the corresponding Fourier-transform *Â*(***k***) looks like the plane-wave exp (−i***k*** · ***x***_0_). The Fourier–Bessel representation 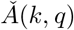 is essentially supported in a cone with ∥*q*∥ = 𝒪(*k*).

**Appendix Figure 5:**
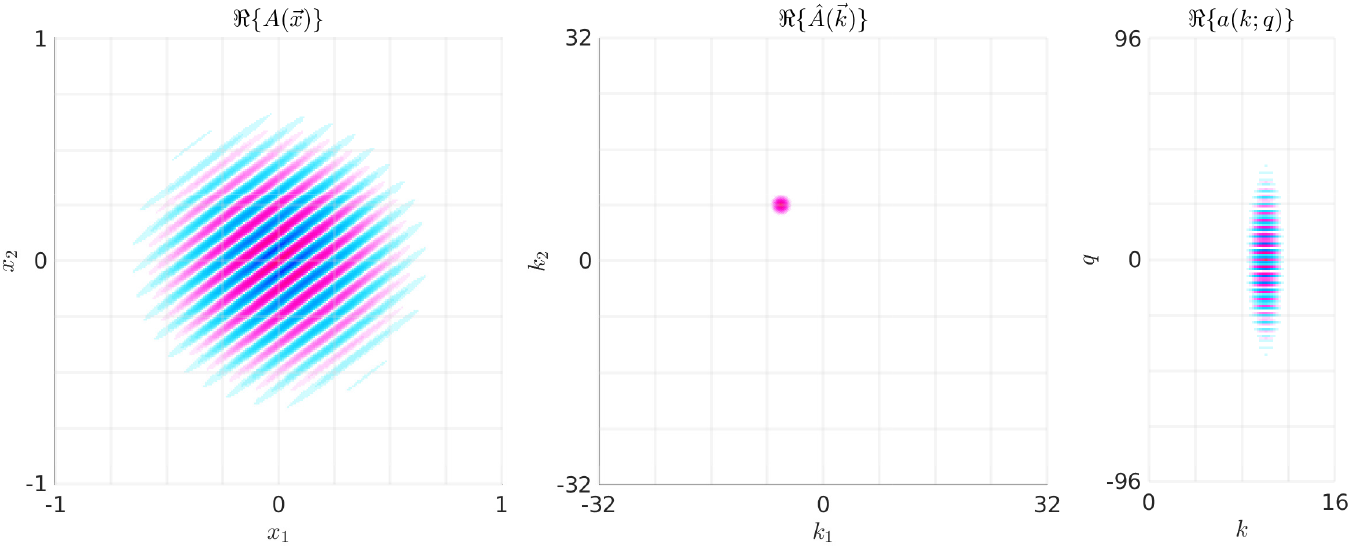
This figure is similar to Fig 4. This time the image has been chosen so that *Â*(***k***) is an isotropic Gaussian in frequency space, with a mean given by ***k***_0_ = [−6, +8]. Thus, *Â*(***k***) is a mollified version of *δ*(***k*** − ***k***_0_). Consequently, the corresponding real-space function *A*(***x***) looks like the plane-wave exp (+i***k***_0_ · ***x***). The Fourier– Bessel representation 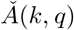 is essentially supported near the vertical strip with *k* = ∥***k***_0_∥ = 10.

**Appendix Figure 6:**
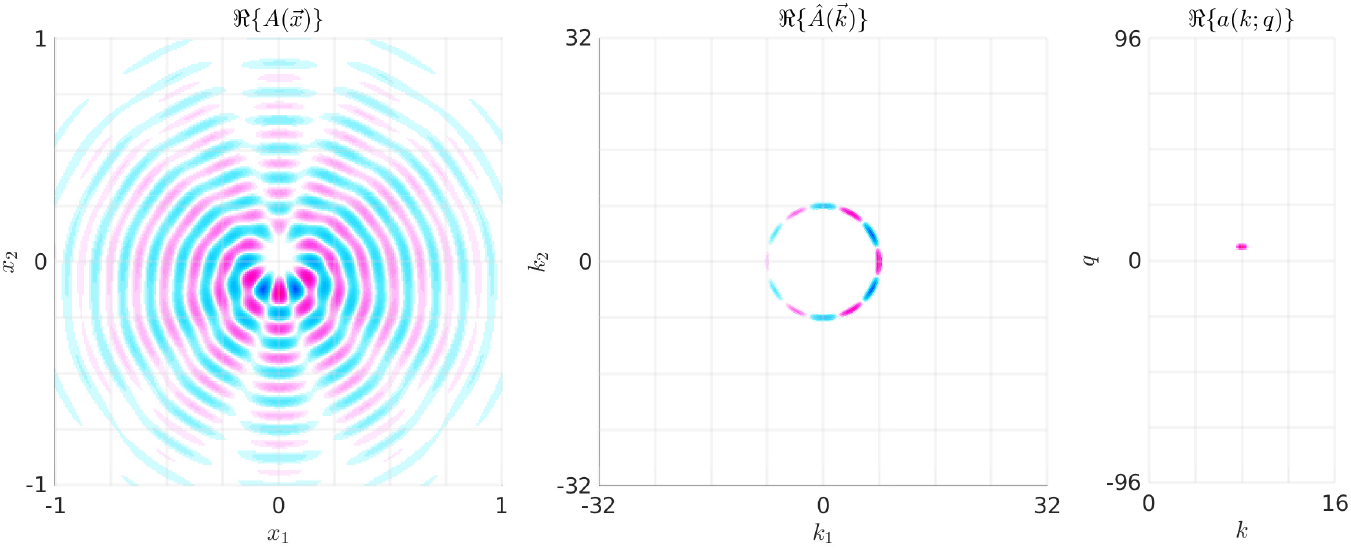
This figure is also similar to fig 4. This time the image has been chosen so that 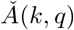 is sharply peaked around *q* = 5 and *k* = 8. Consequently, the corresponding Fourier-space function *Â*(***k***) looks like a ring at *k* = 8 with *q* = 5 oscillations. The real-space function *A*(***x***) looks like a Fourier–Bessel function with a radial component determined by *J*_*q*_(*kx*).

## Appendix Tables

**Appendix Table 1:** List of default CryoLike parameters

| Parameter | Allowed range | Default value |
| --- | --- | --- |
| resolution factor | $(0, 1]$ | 1.0 |
| maximum 2D displacement in pixels | $[0, n_p/2]$ | 16.0 |
| total number of displacements | $[0, \infty)$ | $1024 = 32^2$ |
| atomic models (or radii) | $\{(0, \infty)\}$ | use VDW model [20] |
| viewing angle distance | $[0, 4\pi)$ | $1.0/(4\pi)$ |
| number of in-plane angles | $[0, \infty)$ | 256 |

**Appendix Table 2:** CryoLike parameters used for the wall clock time benchmark (Fig. 2) and for the tests with experimental images (Fig. 3)

|  | Number of |  |  |  |
| --- | --- | --- | --- | --- |
|  | frequencies | inplane angles | viewing angles | displacements |
| Benchmark |  |  |  |  |
| (a) | 33 | 32 | - | 32 |
| (b) | [33..257] | [64..512] | - | 32 |
| (c) | 33 | [64..512] | - | [1..4096] |
| Protein |  |  |  |  |
| apoferritin | 133 | 256 | 2370 | 1024 |
| hemagglutinin | 257 | 256 | 2370 | 4096 |
| spike | 201 | 256 | 2370 | 1024 |

## Appendix Text

### A Mathematical details

In this Appendix, we describe in more detail the computations mentioned within the main text. The software CryoLike is designed to carry out these computations within a Python workflow that allows for GPU acceleration.

We first introduce the notation for our 2D coordinate system in section A.1, as well as our convention for the full-field 2D Fourier transform. In section A.2, we review the rotation- and translation-operators in 2D physical- and frequency-space. In section A.3, we describe the full-field Fourier-Bessel transform, as well as how this basis can facilitate the calculation of in-plane rotations and innerproducts.

The earlier sections above involve 2D functions and their transformations in a continuous setting. The subsequent sections describe the discretization we use for the computations within CryoLike. In section A.6, we describe the polar grid we use to sample the images and templates in frequency-space, as well as the associated bandlimited Fourier-transform. In section A.7 we describe the bandlimited Fourier-Bessel transform, and review how we use this basis to compute bandlimited innerproducts across a range of in-plane rotations.

Up until this point we have only considered 2D functions. The next section A.8 introduces the notation for our 3D coordinate system, as well as our convention for 3D rotations and 3D Fourier transforms. In section A.9 we describe how we discretize the bandlimited frequency-domain in 3D, and how we use the physical representation of the volume to construct 2D templates in frequency-space. In section A.10 we describe how to correct these templates for a point-spread- or contrast-transfer-function, and in section A.11 we describe how we sample the space of possible viewing-angles to accurately resolve the set of all templates.

In section A.12 we move on to the likelihood calculation, starting with a review of our basic image noise-model. From there we describe the basic image-to-2D-template likelihood in section A.13, and the various strategies for marginalization in section A.14. We conclude with a few remarks in section A.15.

#### A.1 Image notation

Within any given image (or template), we use ***x*** ∈ℝ^2^ to represent the position in physical space and ***k*** ∈ ℝ^2^ to represent the position in frequency space. In polar coordinates, these vectors can be represented as:

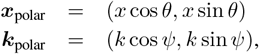

for appropriately chosen angles *θ* and *ψ*.

The full-field (*i.e*., non-bandlimited) Fourier-transform of a two-D function *S* ∈ *L*^2^(ℝ^2^) is defined as

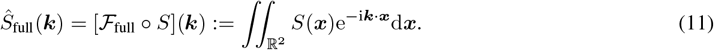

We can recover *S* from 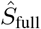 using the inverse Fourier-transform:

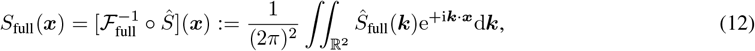

producing an *S*_full_ which is equivalent to the original *S*. The full (*i.e*., non-bandlimited) inner product between two functions *S, A* ∈ *L*^2^(ℝ^2^) is written as

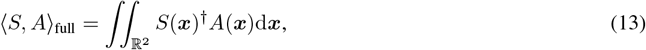

where *z*^*†*^ is the complex conjugate of *z* ∈ ℂ. This full inner product satisfies Plancherel’s theorem [32],

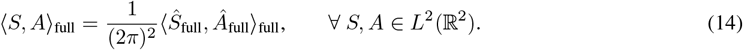

We will ignore the factors of 2*π* associated with the Fourier transform when they are irrelevant.

Below, we will refer to the inner product between an image and itself, which we will denote by the standard l2-norm:

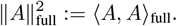

In a slight abuse of notation, we will refer to real-space functions *A*(***x***) and their Fourier-transforms **Â**(***k***) in polar coordinates as:

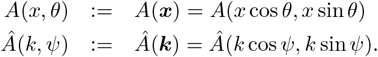

With this notation, each **Â**(*k, ψ*) for fixed *k* and *ψ* ∈ [0, 2*π*) corresponds to a ‘ring’ in frequency space with radius *k*.

#### A.2 Rotation and translation

Using the notation above, an ‘in-plane’ rotation ℛ(*γ*) of an image *A*(***x***) by angle *γ* ∈ [0, 2*π*) can be represented as:

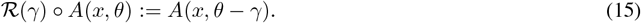

Since rotation commutes with the Fourier-transform, we have:

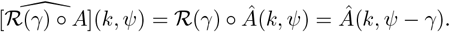

In this manner, a rotation of any image by +*γ* can be represented as an angular shift of each image-ring by *ψ* → *ψ* − *γ*. Likewise, a translation 𝒯 (***δ***) of an image *A* by the shift vectorδ ∈ ℝ ^2^ can be represented as:

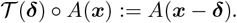

In the Fourier domain, this translation can be expressed as

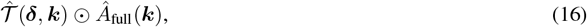

where the translation kernel 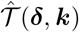 is

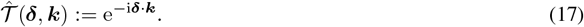

#### A.3 Fourier-Bessel coefficients

Throughout our likelihood calculation, we calculate many image-template inner products across a large collection of (typically equispaced) in-plane rotations *γ*. To ease the computational burden associated with these calculations, we use the Fourier-Bessel basis [22, 23, 33–35].

To define the Fourier-Bessel coefficients of an image, recall that each image-ring **Â**_full_(*k, ψ*) (for fixed *k*) is a 2*π*-periodic function of *ψ*. This image-ring can itself be represented as a Fourier series in *ψ* using the Fourier-Bessel transform B_full_ and its inverse 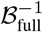:

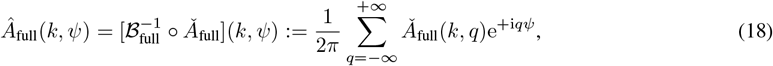

for *q* ∈ 𝕫. The Fourier-Bessel coefficients 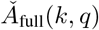 of the image ring **Â**_full_(*k, ψ*) are given by

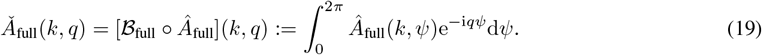

These coefficients can be represented more traditionally by recalling that the Bessel function *J*_*q*_(*kx*) can be written as:

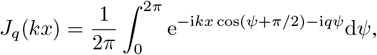

which, when combined with the definition of the Fourier-transform, implies that:

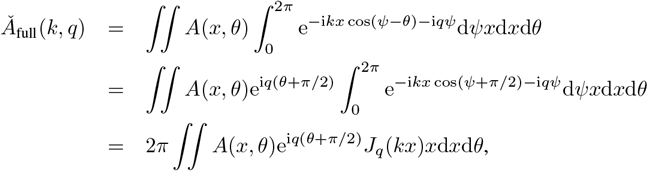

which is the inner product between the original image (in real space) and a scaled Bessel function. Examples of these different representations are given in Appendix Figs 4-6. Note that, using the Fourier-Bessel representation of an image, the rotation ℛ (*γ*) can now be represented as:

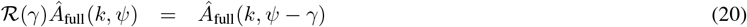

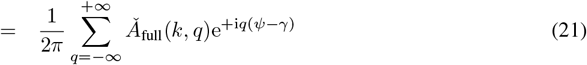

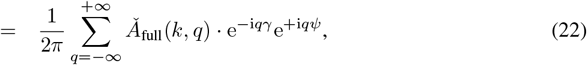

such that the Fourier-Bessel coefficients of the rotated image-ring ℛ ∘ (*γ*) **Â**_full_(*k*,.) are given by the original Fourier-Bessel coefficients 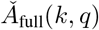, each multiplied by the phase-factor e^−i*qγ*^.

#### A.4 Image inner products in a continuous setting

Note that Eq. (22) naturally allows for arbitrary rotations by any *γ*, requiring only a pointwise multiplication by the appropriate phase factor. This observation allows for the efficient alignment of one image to another (see section A.7 below, and refs. [22, 23, 35]). In practice, we typically apply in-plane rotations in Fourier–Bessel coordinates. For example, using Plancherel’s theorem in 1-dimension, we see that, for any pair of image rings,

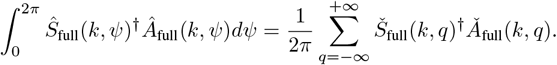

Thus, the inner product

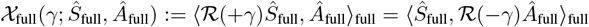

can be written in terms of Fourier–Bessel coefficients of *Ŝ*_full_ and **Â**_full_:

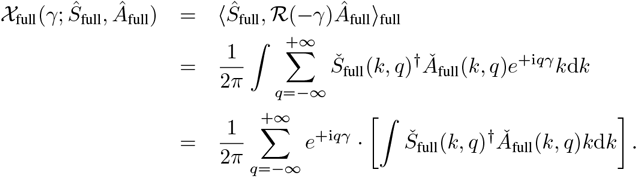

This last expression can be interpreted as a relationship between the desired inner products 𝒳 (*γ*) and the Fourier-Bessel transform of the term in brackets on the right-hand side. That is:

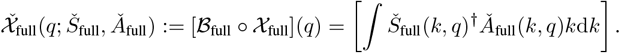

In a typical discretization scheme (see below), the bandlimited images *Ŝ*_band_ and **Â**_band_ each require the storage of 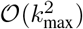 values, as do the Fourier–Bessel representations **Š**_band_ and 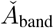. The calculation of the bandlimited array 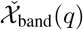 involves 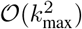 operations and 𝒪 (*k*_max_) storage. Once 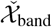 is calculated, the bandlimited inner products 𝒳_band_(*γ*) can be recovered on a uniform grid of 𝒪 (*k*_max_) angles *γ* ∈ [0, 2*π*) with an additional 𝒪 (*k*_max_ log *k*_max_) operations using the Fourier-Bessel transform (i.e., the 1-D fast-Fourier-transform -FFT). More details are given in section A.7 below, and in refs. [22, 23].

#### A.5 Bandlimited approximation

We denote by Ω(*x*_max_) the domain |***x***|_∞_ ≤ *x*_max_, which in 2D corresponds to the unit-square [−*x*_max_, +*x*_max_]^2^. We denote by Ω(*k*_max_) the domain |***k***| ≤ *k*_max_, which in 2D corresponds to the disc of radius *k*_max_. We assume that all the images *A*(***x***) considered are supported in Ω(*x*_max_). The support constraint Supp[*A*] ⊆ Ω(*x*_max_) implies that the representation **Â**(***k***) will itself have a band limit of *x*_max_. This band limit, in turn, implies that **Â** can be accurately reconstructed from its values sampled on a frequency grid with spacing of 𝒪 (1*/x*_max_) [27]. In practice, *x*_max_ is typically several hundred Angstroms. For the calculations in our code, we usually use non-D units in which *x*_max_ is set to 1.

The domains Ω(*x*_max_) and Ω(*k*_max_) motivate our use of the bandlimited Fourier-transform:

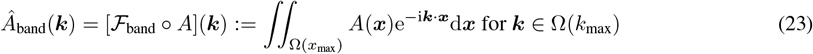

and its bandlimited inverse:

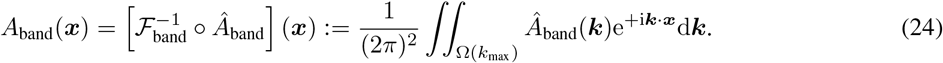

While **Â**_band_ will only approximate **Â**_full_, and *A*_band_ will only approximate the original *A*, we operate under the assumption that these approximations are accurate, at least up to the resolution requested by the user. That is to say, we assume that the signal within the images has a maximum effective spatial-frequency magnitude of *k*_max_; *i.e*., that the features of **Â** we wish to capture are concentrated in ***k***∈ Ω(*k*_max_). Thus, for example, we will only ever compare features of *A*_band_ to features of the original *A* when those features involve frequencies smaller than *k*_max_; *i.e*., we will ignore high-frequency ringing artifacts associated with the band-limiting process.

Given these assumptions, we will use the bandlimited spatial inner product:

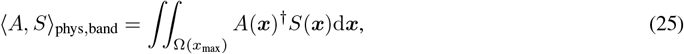

which is identical to ⟨ *A, S* ⟩_full_ so long as *A* and *S* are supported within Ω(*x*_max_). We will also use the bandlimited Fourier inner product:

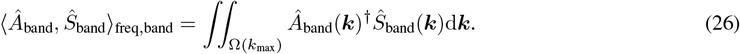

Similar to the bandlimited Fourier-transform above, we operate under the assumption that Plancherel’s theorem is approximately true for these bandlimited inner products, *i.e*.,

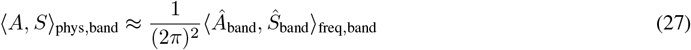

for the images we are considering.

Similar to the full-field scenario, we will often consider bandlimited inner products between an image and itself, which we denote as:

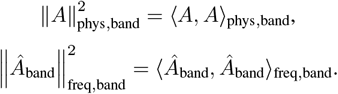

The fact that *x* ≤ *x*_max_ implies that the Bessel coefficients **Š**_full_(*k, q*) will be concentrated in the range |*q*| ≲ *kx*_max_, meaning that the Bessel coefficients **Š**_full_(*k, q*) across all *k* ∈ [0, *k*_max_] will be concentrated in *q* ∈ [−*q*_max_*/*2, +*q*_max_*/*2 − 1] for *q*_max_ = 𝒪 (*k*_max_*x*_max_) = 𝒪 (*N*_pix_*π/*2). Thus, we can use the bandlimited Fourier-Bessel-transform:

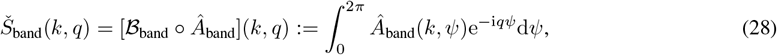

for *q* ∈ [−*q*_max_*/*2, +*q*_max_*/*2 − 1], and and

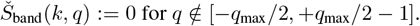

its bandlimited inverse:

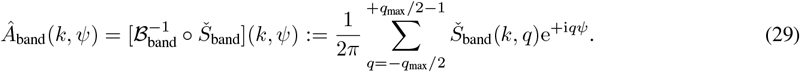

#### A.6 Image discretization

With the notation above, we can consider an *n*_p_ × *n*_p_ image as a discrete set of pixel-averaged samples within [−*x*_max_, +*x*_max_]^2^:

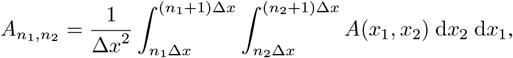

for indices *n*_1_, *n*_2_ ∈{0, …, *n*_p_ − 1}. We approximate the (bandlimited) Fourier-transform **Â**_band_ at any ***k*** Ω(*k*_max_) via the simple summation:

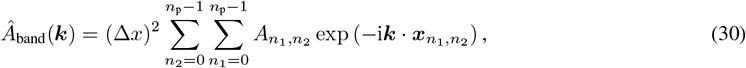

where 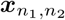 is the appropriately-chosen pixel-center (e.g., 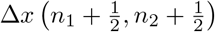). Note that the largest *k*_max_ that we can reasonably expect to attain is related to the number of pixels *n*_p_ along a side in the original image. More specifically, if we assume that *A*(***x***) is supported in ***x*** ∈ Ω(*x*_max_), then the pixel-spacing will be Δ*x* = 2*x*_max_*/n*_p_. The alignment of the pixel-grid to the support then implies that the Nyquist spatial-frequency is given by *π/*Δ*x* = *n*_p_*π/*(2*x*_max_). For many typical applications *k*_max_ will be smaller than *n*_p_*π/*(2*x*_max_), particularly when the goal is to analyze low-resolution features of images and volumes.

Because we have assumed that the image is sufficiently well sampled (*i.e*., that **Â** contains little relevant frequency content above the Nyquist frequency), we expect the simple sum above to be accurate. However, we remark here that, while ℛ (*γ*) commutes with both the full- and bandlimited-Fourier-transform, the same does not hold for 𝒯 (***δ***). Indeed, if 𝒯***δ*** is sufficiently large that (***δ***) shifts the support of *A*(***x***) outside the domain Ω(*x*_max_), then we expect relevant information to be lost when applying the bandlimited operators above. Thus, in our calculations, we will always assume that the maximum in-plane translation δ_max_ is sufficiently small that this support constraint is not violated. That is, we will assume thatδ_max_ is sufficiently small that ℛ (*γ*) 𝒯 (***δ*** s.t. |δ |***δ***_max_) ℛ (*γ*^*′*^)*A*(***x***) remains supported within Ω(*x*_max_) for all choices of *γ* and *γ*^*′*^. Equivalently, if a largeδ_max_ is desired, one can increase *x*_max_ to accommodate the extended support associated with such a large translation. This increase in *x*_max_ will then mandate an increase in *k*_max_ to remain at the Nyquist sampling frequency (as the pixel grid for the original images is no longer aligned with the extended support).

We will typically evaluate **Â**_band_(***k***) for ***k*** on a polar-grid, with *k*- and *ψ*-values corresponding to suitable quadrature-nodes, using the 2D non-uniform FFT (2D NUFFT) to compute **Â**_band_(*k, ψ*) at those nodes (see refs. [23, 27]). The quadrature in *k* corresponds to a set of *n*_*k*_ nodes: 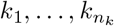 and radial weights 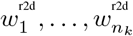 so that

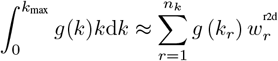

to high accuracy for a smooth function *g*(*k*). The exact radial quadrature-nodes we use correspond to a Gauss-Jacobi quadrature for *k* built with a weight-function corresponding to a radial weighting of either *k*d*k* (for images) or *k*^2^d*k* (for volumes). In both cases we ensure that both the 2- and 3-D integrals are calculated to single-precision; the resulting number of radial quadrature-nodes *n*_*k*_ will be of the order 𝒪 (*k*_max_).

The *q*_max_ angular-nodes 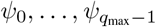 are equispaced in the periodic interval [0, 2*π*), with a spacing of Δ*ψ* = 2*π/q*_max_, and *ψ*_*q*_^′^ = *q*^*′*^Δ*ψ*. These equispaced *ψ*-nodes allow for spectrally accurate trapezoidal quadrature in the *ψ*-direction, and we approximate the bandlimited Fourier-Bessel coefficients of each image-ring **Â**_band_(*k, ψ*) as follows:

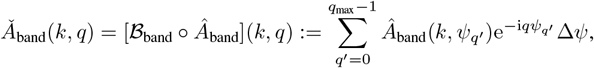

with the index *q* considered periodically in the interval [− *q*_max_*/*2, …, +*q*_max_*/*2 − 1] (so that, e.g., the *q*-value of *q*_max_ 1 corresponds to the *q*-value of 1). Because *q*_max_ is equal to the number of equispaced *ψ*-values, this implementation of the band-limited Fourier-Bessel transform can be applied and inverted using the standard 1-D FFT:

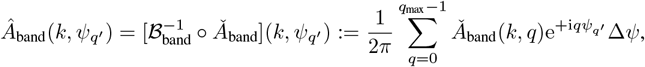

with 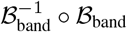 corresponding to the identity. This property allows us to easily transform back and forth between **Â**_band_(*k, ψ*_*q*_^′^) and 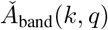 as many times as needed, without incurring any significant loss of accuracy.

#### A.7 Bandlimited image inner products in a discrete setting

As alluded to in section A.4, we can use the discrete Fourier–Bessel coefficients **Š**_band_(*k, q*) and 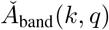 associated with images *Ŝ*_band_ and **Â**_band_ to efficiently calculate the inner products

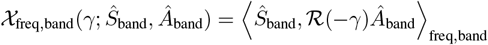

across a range of rotation angles *γ*_*q*_^′^ = 2*πq*^*′*^*/q*_max_. This calculation can be summarized as:

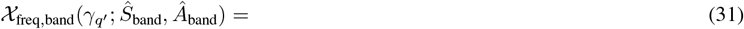

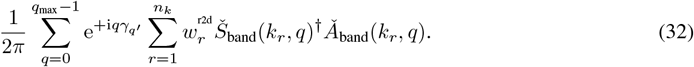

This calculation can be achieved in two steps:

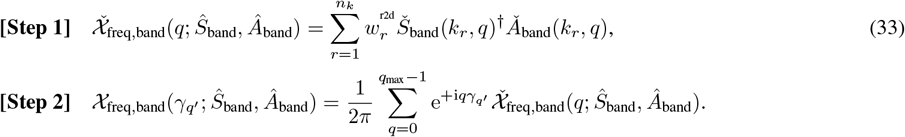

The first step applies radial quadrature to combine information from different image rings, computing the Fourier-Bessel series coefficients 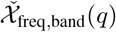 in 𝒪 (*n*_*k*_*q*_max_) operations. The second step can be evaluated using a 1-D FFT of size *q*_max_, requiring 𝒪 (*q*_max_ log(*q*_max_)) operations. Because *n*_*k*_ and *q*_max_ are both 𝒪 (*k*_max_), this entire operation is 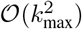 If a finer resolution in *γ* is required then the array 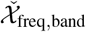 can be zero-padded prior to the 1D FFT. If a specific nonuniform set of *γ*-values are requested, then the 1D NUFFT can be applied instead.

#### A.8 Volume notation and discretization

Within any given volume, we use **x, k** ∈ℝ^3^ to represent spatial position and frequency, respectively. We represent any given volume as a function *F* (**x**) ∈*L*^2^(ℝ^3^), with values corresponding to the volume intensity at each location **x** ∈ [− *x*_max_, +*x*_max_]^3^. Abusing notation slightly, we will characterize this spatial-domain as |**x**| _∞_ ≤ *x*_max_, and (just as in 2D) denote it by Ω(*x*_max_). We will also define the 3-D full-field (non-bandlimited) Fourier-transform (and its inverse) similar to above:

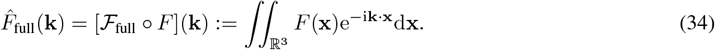

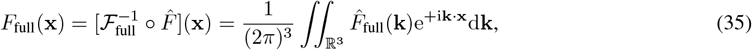

producing an *F*_full_ which is equivalent to the original *F*. Given a resolution-limit of *k*_max_, we define the domain Ω(*k*_max_) to be the ball |**k**|_2_ ≤ *k*_max_, and the corresponding bandlimited Fourier-transform as:

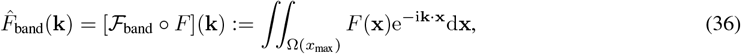

for **k** ∈ Ω(*k*_max_), and

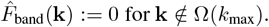

In principle, if we have a density-map *F* (**x**) sampled on an *n*_p_ × *n*_p_ × *n*_p_ grid of voxels describing Ω(*x*_max_), we could approximate the bandlimited Fourier-transform 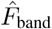 via:

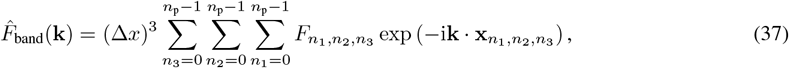

where 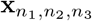 is the appropriately-chosen pixel-center, e.g., 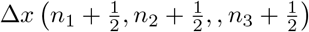. However, as we will discuss in the next section, we do not compute 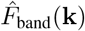 using this standard 3D FFT.

#### A.9 Constructing 2D templates

In spherical coordinates the vector **k** is represented as:

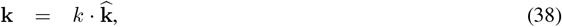

with *k* denoting the amplitude |**k**| and 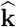 denoting a unit-vector lying on the surface of the sphere *S*^2^. We will refer to 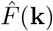 in spherical coordinates as 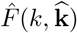.With this notation, each 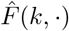 corresponds to a ‘shell’ in frequency-space with radius *k*. Using a right-handed basis, a rotation about the third axis by an azimuthal angle *α* or an in-plane angle *γ* is represented as:

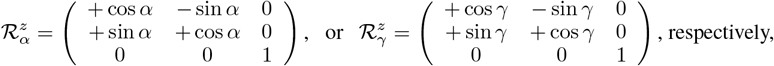

and a rotation about the second axis by angle *β* is represented as:

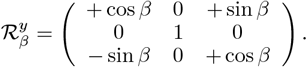

A rotation ℛ (*τ*) of a vector **k** ∈ℝ^3^ can be represented by the vector of Euler angles *τ* = (*γ, β, α*) (for notation simplicity we have omitted the translation from the pose *τ*):

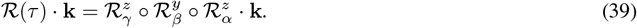

Below we will often deal with scenarios where *γ* = 0, in which case we will refer to *τ* simply by (*β, α*). With this compact notation we will refer to the rotation ℛ(*τ*) simply as ℛ(*β, α*).

In principle, the rotation of any volume 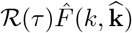 corresponds to the function 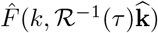.Such a rotation can be cumbersome to evaluate without the appropriate choice of basis, however, in our code we do not compute rotations of this kind. Rather, we take advantage of the Fourier-slice theorem, as decribed below.

To begin with, we denote the full-field (*i.e*., non-bandlimited) projection of a volume *F* (**x**) (with **x** ∈ ℝ^3^) along a particular viewing-angle *τ* = (*γ, β, α*) by the function *S*_full_(***x***; *τ* ; *F*) (with ***x*** ∈ ℝ^2^). We will refer to this projection *S*_full_(***x***; *τ* ; *F*) as the ‘template’ associated with *F* (**x**) and viewing-angle *τ*. By the Fourier-slice theorem, we can decribe the Fourier-transformed template 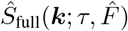 via:

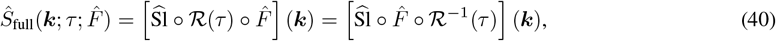

where ***k*** ∈ ℝ^2^ (note that it is different from **k** see below), and 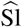 denotes taking the equatorial slice in the Fourier domain. In terms of computation, the 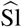 operator can be described by restricting the unit-vector in Eq. 38 to the equatorial-plane, corresponding to setting ***k***_1_ := **k**_1_ and ***k***_2_ := **k**_2_ and ignoring **k**_3_.

Now let’s consider a particular ***k*** ∈ ℝ^2^, and the associated-value of 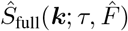.By using the far right-hand-side of Eq. 40, we can see that 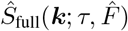 has the same value as 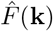,where ℝ(*τ*) ? **k** = [***k***_1_, ***k***_2_, 0]^⊺^. This same relationship holds for every ***k*** ∈ ℝ^2^. Thus, the values of the template *Ŝ*_full_ can also be written as:

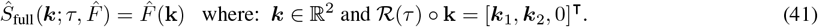

Intuitively, **k** and ***k*** are related via:

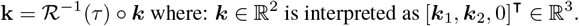

Using this description, we can easily generate the set of **k**-values **K**(*τ*) associated with the bandlimited template 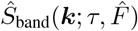.This set **K**(*τ*) is obtained by first defining the set **K**(**0**) of points in the equatorial-plane corresponding to the polar-grid on which we evaluate our bandlimited Fourier-transformed images,

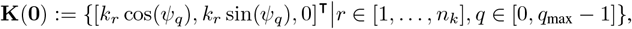

and then rotating this set in 3-D so that it aligns with the location of the desired slice:

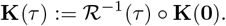

The values of 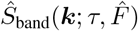 can then be determined directly from the discrete samples 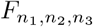 on the voxel-grid by using a 3D NUFFT. To minimize computational overhead we can use the 3D NUFFT to generate the values of multiple templates (*i.e*., involving multiple different viewing-angles *τ*) simultaneously.

#### A.10 PSF and CTF correction

If, furthermore, we wish to consider the effect of a point-spread-function PSF(***x***), then the template associated with *F* (**x**) and a viewing-angle *τ* can be written as:

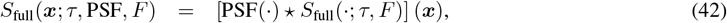

where ‘⋆’ corresponds to a real-space convolution. We will approximate this PSF-correction in Fourier-space with the analogous bandlimited operation:

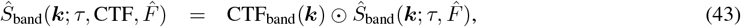

where CTF_band_(***k***) is the bandlimited Fourier-transform of PSF(***x***). These CTF-corrected templates can be calculated by using the strategy above to calculate 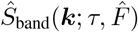,then applying a pointwise-multiplication.

#### A.11 Discretizing the viewing-angles

In practice, we generate the templates described in Eq. 43 for a discrete collection of viewing angles *τ*_*t*_ := (0, *β*_*t*_, *α*_*t*_), for *t* ∈ {1, …, *n*_template_}. As mentioned above, for brevity, we will often refer to *τ*_*t*_ simply by *τ*_*t*_ = (*β*_*t*_, *α*_*t*_). Similarly, we will refer to the rotation ℛ(*τ*_*t*_) as ℛ(*β*_*t*_, *α*_*t*_), the slice **K**(*τ*_*t*_) as **K**(*β*_*t*_, *α*_*t*_), the template 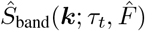 as 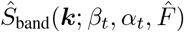,and so forth. To define the {*τ*_*t*_}, we first design a quadrature-scheme on a spherical-shell of radius *k*_max_ (*i.e*., the surface of Ω(*k*_max_)). We design this quadrature scheme so that it can accurately calculate surface integrals of any function 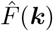 with a bandlimit of *x*_max_. One convenient choice is to select unique values of cos(*β*) to align with Legendre-nodes on [−1, +1], and then for each unique polar-angle *β* we select sufficiently many equispaced azimuthal-angles *α* to accurately resolve that latitudinal-line given the band limit of *x*_max_. This process produces 𝒪(*k*_max_) unique polar-angles, and 𝒪(*k*_max_) azimuthal-angles for each such polar-angle, for a total of 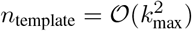 quadrature-nodes. Once we have designed the quadrature for this surface-integral, we define the collection {(*α*_*t*_, *β*_*t*_)} to align with these *n*_template_ quadrature-nodes and save the quadrature-weights 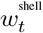 for later use (see Eq. 73 later on).

Note that the particular choice of the third Euler angle (*γ*) for each of the templates is dictated by the orientation of the polar grid used in **K**(**0**). Since we expect our polar grid to be sampled accurately in the angular direction, and we expect to consider each template across a well-sampled grid of in-plane rotations, this third Euler angle does not play an important role in our calculations. More specifically, for *τ* = (*γ, β, α*) we have that:

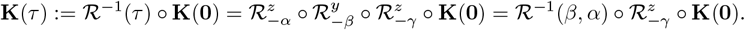

This immediately implies that:

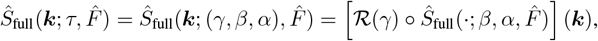

where the operator ℛ (*γ*) on the right-hand-side corresponds to a 2D rotation by *γ* in image-space. A similar statement holds for the CTF- and PSF-corrected templates in both full- and bandlimited-form.

#### A.12 Image intensity and noise model

We assume that the images considered can be modeled as the sum of a ‘signal’ plus a ‘noise’. We assume that the signal corresponds to a 2-D projection *S*_full_(***x***; *τ* ; PSF; *F*) of a (smooth) molecular density function *F* (**x**), while the noise corresponds to detector noise [2, 36–38]. We do not consider more complicated sources of noise, such as structural noise associated with image preparation [39].

Based on these simple assumptions, we will model the noise in real space as independent and identically distributed (iid), with a (possibly image-specific) variance of *λ*^2^ at each pixel in real space. Thus, we expect a typical discretely-sampled image-array 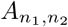 from that molecule to take the form:

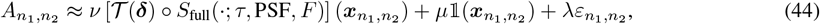

where *τ*, ***δ*** and PSF are the viewing-angle, displacement and point-spread-function associated with the imaging-process, *ν* is an image-specific intensity factor, *µ* is an image-specific offset, 𝟙(***x***) is the indicator-function for the spatial-domain Ω(*x*_max_) and *λ* is the standard deviation of the noise. The discretely sampled noise array is denoted by 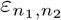,and we assume that each pixel of this noise array is drawn independently from N(0, 1).

#### A.13 Image-to-2D-template likelihood

As a direct corollary of Eq. 44, we can calculate the probability that the discretely-sampled image-array 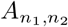 is observed, given the molecule *F* and the point-spread-function PSF, as well as the various parameters *λ, ν, µ*, ***δ***, *τ* :

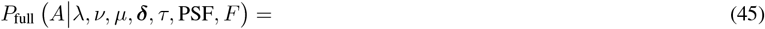

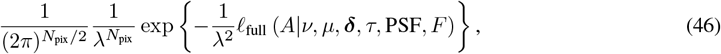

with:

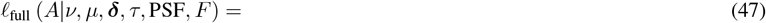

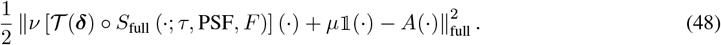

The quantity ℓ is closely related to the log-likelihood of observing the image, given the model and its parameters. In practice, we will assume that the full likelihood in Eq. 46 can be approximated by its bandlimited real-space version:

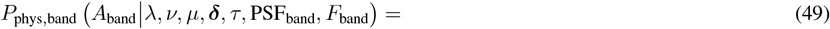

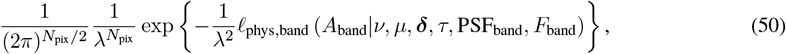

with:

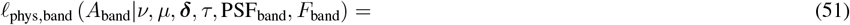

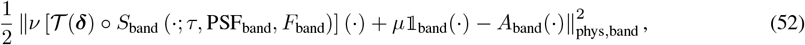

and its bandlimited frequency-space version:

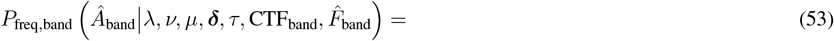

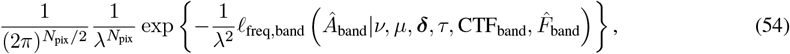

with:

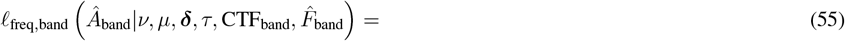

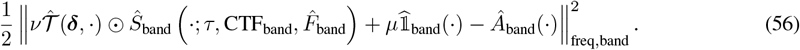

Note that ℓ is a simple function of the image and template. For example, given a particular template *τ*_*t*_ = (*β*_*t*_, *α*_*t*_) and in-plane rotation *γ*, the ℓ_freq,band_ can be decomposed as follows:

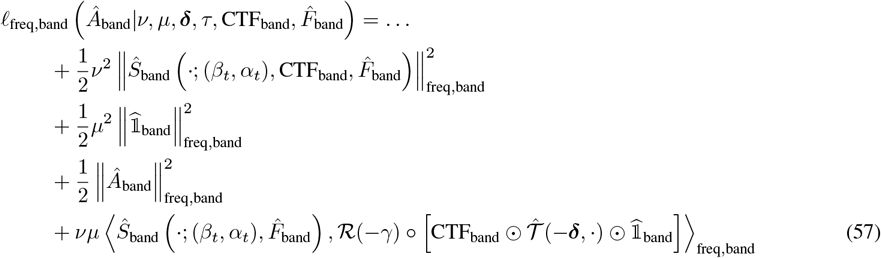

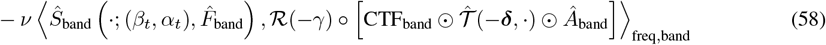

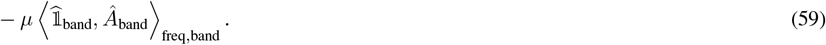

Note that the bandlimited inner products in Eqs 57,58 can be calculated using the strategy outlined in section A.7.

#### A.14 Marginalizing the likelihood

In this section, we describe some general strategies for marginalizing the basic likelihood presented above. For clarity of presentation we will discuss the frequency-space calculations described in Eqs. 54 and 56. To ease notation below, we will drop the subscripts of ‘freq’ and ‘band’.

##### marginalizing with respect to *ν* and *µ*

The likelihoods presented in the previous section take the form of:

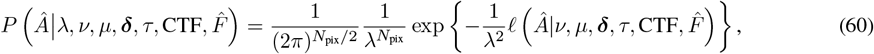

with:

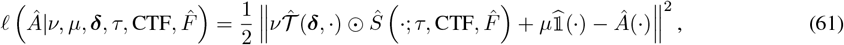

We first marginalize this likelihood with respect to the image-intensity *ν* and/or image-offset *µ* by assuming a uniform prior. We can use Gradshteyn and Ryzhik 2.33:

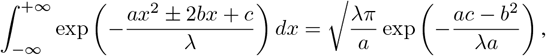

which holds for any normalizing factor *λ*.

Using this relationship, we can write:

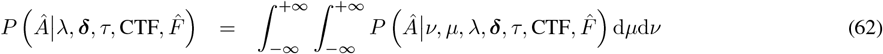

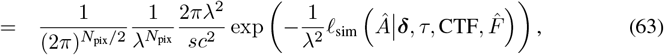

with:

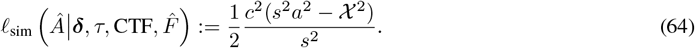

In this expression we denote by *c*^2^ the (bandlimited) *l*2-norm of the indicator-function] 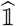:

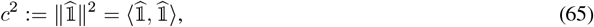

where we expect 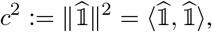. Similarly, we denote the template via:

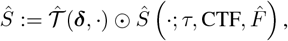

and then calculate the averate template value:

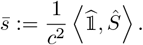

The average template-value can then be used to define the template-variance:

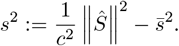

Similarly, we can define the average image-value:

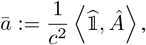

and the image-variance:

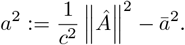

The quantity Χ represents the image-template covariance, defined as:

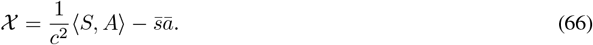

Note that ℓ_sim_ can be directly interpreted as a measure of image-template similarity. Indeed, if the image and template are both normalized to have means 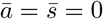 and variances of *a*^2^ = *s*^2^ = 1, then

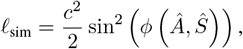

where *ϕ*(**Â*, Ŝ*) is the ‘angle’ between the image and template when they are both considered as vectors in the appropriate space (e.g., 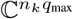).

##### marginalizing with respect to *λ*

The basic likelihood considered in Eq. 54 as well as the marginalized version derived from Eq. 63 both have the general form:

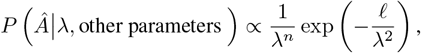

for some number of degrees of freedom *n* and some rate-function ℓ.

Now, assuming a uniform prior over *λ*, we can use Gradshteyn and Ryzhik 3.326.2: and see that:

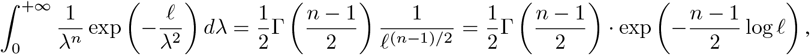

and see that:

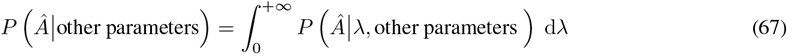

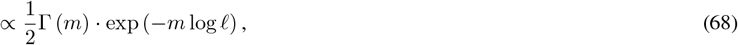

for *m* = (*n* − 1)*/*2.

For the specific case shown in Eq. 63, a marginalization with respect to *λ* produces:

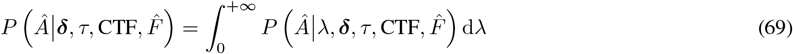

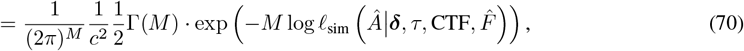

with *M* = (*N*_pix_ − 3)*/*2, and

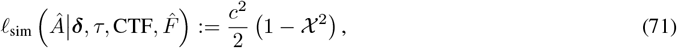

where we have assumed that both the image and template have been normalized to have mean 0 and variance 1, and used Eqs. 65 and 66 to define *c*^2^ and Χ.

##### marginalizing with respect to *τ* and *δ*

The likelihoods presented above are of the general form:

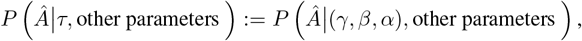

for the viewing-angle *τ* := (*γ, β, α*). We can marginalize with respect to *τ* by using our discrete viewing-angles {(*α*_*t*_, *β*_*t*_)} and the associated quadrature-weights 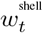 defined in section A.11. This discretization produces:

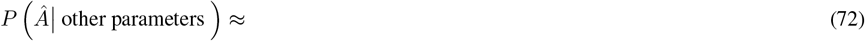

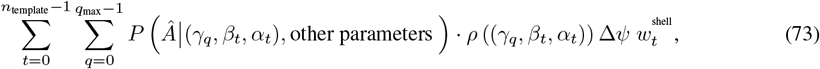

where *ρ*(*τ*) is a prior distribution for the viewing angles *τ* ∈ *SO*3 (which is itself assumed to be well-resolved by our discretization of *SO*3).

A very similar strategy can be used to marginalize with respect to ***δ***, given a set of discrete displacements and associated quadrature weights. Note that, when marginalizing Eq. 70 over *τ* and ***δ***, the value of *M* is often quite large, and thus this marginalized likelihood is typically very well approximated using standard asymptotics (e.g., Laplace’s approximation).

#### A.15 Application to numerical experiments

In the main text we denote by {*τ}* the collection {*τ}* = *γ, β, α*, ***δ***. This *τ* encapsulates the pose parameters for a particular image. In the numerical examples presented in the main text, we consider the ‘maximum-likelihood’ pose *τ* ^opt^ associated with image **Â**, determined via:

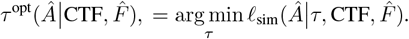

Note that this optimal-pose does <u>not</u> depend on *µ* or *ν*, as these have been marginalized to produce ℓ_sim_. Instead, given our assumption that the images and templates are all normalized to have means of 0 and variances of 1, the ℓ_sim_ only depends *χ* on. Thus, the optimal *τ* is the one that maximizes the image-template covariance *χ*, which is equivalent to the cross-correlation discussed in the main text.

As a technical detail, we can readily calculate the bandlimited frequency-space cross-correlations *χ*_freq,band_ between **Â**_band_ and any particular template *Ŝ*_band_ using Eq. 26 and the strategy described in section A.7. However, calculating the analogous bandlimited physical-space cross-correlations *χ*_phys,band_ is not as easy, as we must first calculate the bandlimited real-space functions *A*_band_ and *S*_band_ via Eq. 24 before we calculate their innerproduct via Eq. 25. For this reason we only calculate *χ*_phys,band_ for *ϕ*^opt^, and refrain from calculating *χ*_phys,band_ across the full collection of *ϕ*.

Because we have access to the bandlimited frequency-space calculations across the full collection of *ϕ*, we have the luxury of marginalizing the bandlimited frequency-space likelihoods with respect to the viewing-angles and displacements:

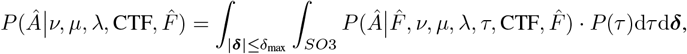

as described by the numerical approximation in Eq. 73 above.

